# IL-13/IL-13Rα2 axis promotes proliferation of angiosarcoma cells

**DOI:** 10.1101/2024.10.24.619789

**Authors:** Hinako Saito, Issei Omori, Okuto Iwasawa, Ayaka Sugimori, Hibari Nakajima, Ryuzo Ichimura, Shinichi Sato, Hayakazu Sumida

**Affiliations:** Department of Dermatology, Graduate School of Medicine, The University of Tokyo, Tokyo, Japan; Scleroderma Center, The University of Tokyo Hospital, Tokyo, Japan; SLE Center, The University of Tokyo Hospital, Tokyo, Japan

**Keywords:** Angiosarcoma, Interleukin-13, IL-13 receptor alpha2, Cell proliferation, STAT6, Cytokine signaling

## Abstract

Angiosarcoma is a rare and aggressive soft tissue sarcoma with a poor prognosis and limited treatment options. The role of interleukin-13 (IL-13) and its receptors in angiosarcoma pathogenesis has been largely unknown. We detected high IL-13 receptor α2 (IL-13Rα2) expression in angiosarcoma cell lines and patient samples compared to other cell types and benign vascular tumors. Moreover, histological analysis showed the presence of IL-13 in the angiosarcoma microenvironment. Functional studies using angiosarcoma cell lines, MO-LAS-B cells, revealed the promoting effect of IL-13 on cell proliferation. The effect was inhibited by siRNA-mediated knockdown of *IL13RA2* or neutralizing antibodies against IL-13, suggesting the direct impact of IL-13/IL-13Rα2 axis in the angiosarcoma proliferation. In addition, IL-13 stimulation increased mRNA levels of *IL13RA2* and *VEGFA*, suggesting an underlying positive feedback mechanism, which was attenuated by a STAT6 inhibitor. These findings highlight the importance of the IL-13/IL-13Rα2 axis in angiosarcoma progression and its potential as a novel therapeutic target for this challenging malignancy.

## Introduction

Angiosarcoma is a rare and aggressive soft tissue sarcoma of vascular or lymphatic endothelial origin^1^. Angiosarcoma frequently presents on the skin surface but also develops in the breast, vascular system, or internal organs. Head and neck are the most common sites for cutaneous angiosarcoma (59.3%), with the scalp being most frequently affected^1,2^. Cutaneous angiosarcoma predominantly affects patients over 60 years of age, accounting for approximately 85% of cases^3^. The 5-year survival rate for cutaneous angiosarcoma ranges from 38.6% to 45%, which is the lowest among cutaneous soft tissue sarcoma, compared to the 5-year survival rates of 99%, 89%, and 92% for dermatofibrosarcoma protuberans, malignant fibrous histiocytoma, and leiomyosarcoma, respectively^1,3^. The prognosis for angiosarcoma is typically poor, characterized by infiltrative growth and high metastatic potential, which significantly increases the risk of recurrence and metastasis^1,3^. Currently, therapeutic strategies include complete surgical resection with negative margins, often accompanied by preoperative or postoperative radiation therapy. For metastatic or unresectable cases, anthracycline-based or taxane-based chemotherapy regimens are frequently employed^4^. However, the efficacy of these treatments remains limited in most patients^5^.

Regarding the pathogenesis of angiosarcoma, vascular endothelial growth factor (VEGF) is the most well-known factor involved in proliferation, while other factors such as interleukin-6 (IL-6) have also been identified^6^. VEGF is a potent angiogenic factor upregulated in many malignancies, crucial for tumor angiogenesis and survival. Inhibiting the VEGF-A/VEGFR2 pathway suppresses angiogenesis and tumor growth^7^. VEGF-A and VEGFR-2 are expressed in angiosarcoma cells, correlating with proliferation^8^, suggesting that VEGF-targeting drugs may be promising for angiosarcoma treatment. Anti-angiogenic therapies, including VEGF-R inhibitors (sorafenib, pazopanib) and anti-VEGF antibodies (bevacizumab), have been considered, though initial studies show limited efficacy as monotherapies^9^.

Recent studies have focused on interleukin-13 (IL-13) and its receptor interleukin-13 receptor a2 (IL-13Ra2) in malignant tumor progression across various cancers, including melanoma, renal cell carcinoma, adrenocortical carcinoma, and brain tumors^10–13^. Overexpression of IL-13Rα2 has been correlated with advanced disease and poor prognosis in colorectal, gastric, lung, and renal cancers, as well as in glioblastoma multiforme^14–19^. In terms of IL-13 signaling, IL-13 is known to have two receptors, IL-13Rα1 and IL-13Rα2. IL-13Rα1 forms a heterodimer with interleukin-4 receptor α (IL-4Rα) and drives a JAK-dependent phosphorylation of the transcription factor STAT6^20,21^. On the other hand, IL-13Rα2 has a higher affinity for IL-13 than IL-13Rα1^22^. Structural analysis revealed that IL-13Rα2 lacks the conserved JAK binding sites essential for signal transduction, in contrast to IL-13Rα1 and IL-4Rα. Consequently, IL-13Rα2 has long been considered a non-signaling decoy receptor in the interleukin-13 signaling pathway^23^. However, recent evidence demonstrates that IL-13 can effectively signal through IL-13Rα2 in human cells. This signaling occurs via STAT6-dependent or independent pathways involved in the pathogenesis of atopic dermatitis or tumor progression^11,18,24,25^. Although whole genome sequencing analysis of angiosarcoma has suggested the possibility of increased expression of IL-13Rα2 in angiosarcoma^24^, to the best of our knowledge, there have been no studies focused on the pathological roles of IL-13Rα2 in angiosarcoma. Therefore, here we decided to investigate these aspects using clinical samples and angiosarcoma cell lines.

## Results

### High expression of IL-13Rα2 in angiosarcoma cells with the presence of IL-13

Given the difficulty in isolating primary angiosarcoma cells from patients, we first examined the mRNA expression of IL-13 receptors using recently established and reported angiosarcoma cell lines ISO-HAS-B and MO-LAS-B to investigate the effect of IL-13 on angiosarcoma. For comparison, we sorted various immune cells from peripheral blood and cultured primary human skin fibroblasts from healthy donors. We further prepared human skin squamous cell carcinoma cell line HSC5 and human epidermoid carcinoma cell line A431. Our quantitative reverse-transcription PCR (QPCR) analysis confirmed high expression of IL-13Rα1 in peripheral blood neutrophils (Fig. 1A), which is consistent with the Human Protein Atlas database (www.proteinatlas.org) and the mouse study^26^. Importantly, our findings revealed that angiosarcoma cell lines exhibited much lower IL-13Rα1 expression compared to neutrophils and other peripheral T and B cells (Fig. 1A). On the other hand, we observed markedly higher IL-13Rα2 expression in angiosarcoma cell lines relative to any other examined cell types (Fig. 1B). Further flow cytometry analysis confirmed these findings at the protein level, revealing higher IL-13Rα2 signals in angiosarcoma cells compared to A431 cells (Fig. 1C), consistent with our QPCR results (Fig. 1B). To verify our findings in clinical samples, we next performed section staining on patient-derived angiosarcoma samples to examine the protein expression levels of IL-13Rα2. Strong signals were observed in angiosarcoma cells (Fig. 1D), while weak or no signals were detected in hemangioma, a benign vascular tumor (Fig. 1E). These findings from immunohistochemical staining support our QPCR results (Fig. 1B).

**Fig 1.**
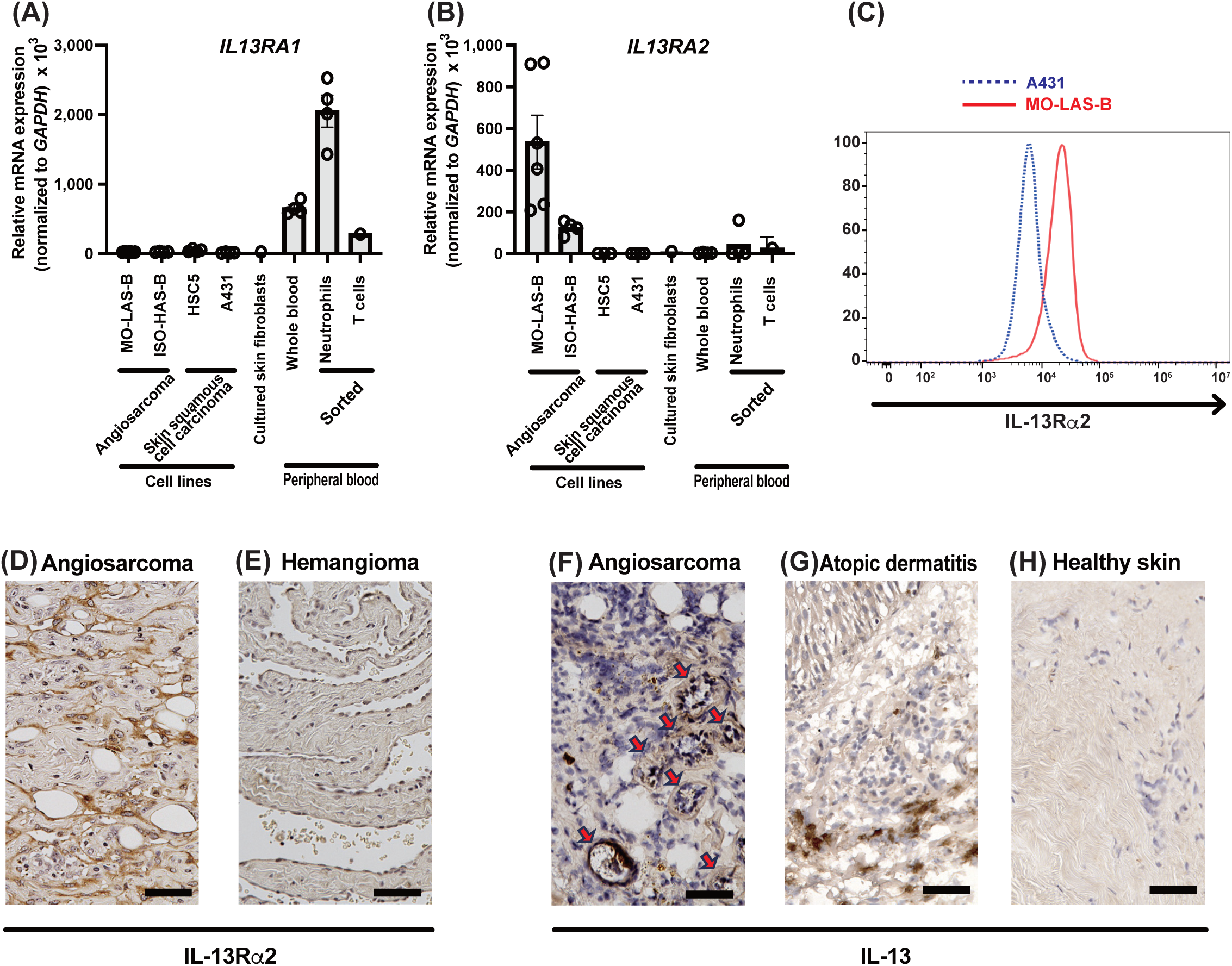
IL-13 and IL-13 receptors expression in angiosarcoma and control cells. (A) Relative mRNA expression of *IL13RA1* (A) and *IL13RA2* (B) in various cell types. (C) Surface protein expression levels of IL-13Ra2 in MO-LAS-B and A431 cells. Representative images from immunohistochemical staining for IL-13Ra2 (D, E) and IL-13 (F-H) in angiosarcoma (D, F), hemangioma (E), atopic dermatitis (G), and healthy skin (H) samples. A subset of atypical tumor cells forming luminal structures (red arrows). Scale bars = 50μm.

On a related note, we also considered it necessary to confirm the abundance of the IL-13Rα2 receptor ligand, IL-13 in angiosarcoma tissue. Immunolabelling for IL-13 showed strong signals in a subset of tumor cells from angiosarcoma patients, particularly in atypical tumor cells forming luminal structures (Fig. 1F). Since IL-13 is known to be expressed in infiltrated lymphocytes to the dermis in atopic dermatitis^27^, human lesional samples from atopic dermatitis were used as a positive control, which showed positive staining only in lymphocyte-like cells in the upper dermis (Fig. 1G). In parallel, samples from healthy skin exhibited no signals (Fig. 1H). These histological observations indicate that IL-13 exists in the environment surrounding angiosarcoma cells. Based on these expression analyses, we proceeded to investigate the effects of IL-13 and its receptors in angiosarcoma to further characterize their functional significance using MO-LAS-B cells which showed the highest expression of IL-13Rα2 in our QPCR analysis.

### Promoted proliferation of angiosarcoma cells mediated by IL-13/IL-13Rα2 axis

We next investigated the effect of IL-13 on angiosarcoma proliferation by experiments with IL-13 supplementation. The IL-13-treated group exhibited increased cell number compared to the vehicle-treated group by two types of evaluation methods (Figs. 2A-B). Further flow cytometry experiments showed that IL-13 treatment (10 ng/mL, 48 hours) significantly increased the percentage of both Ki-67^high^ and BrdU-positive cells compared to vehicle treatment (Figs. 2C-F), indicating the proliferative effect of IL-13 contributed to the increased number of cells under the IL-13 treatment (Figs. 2A-B). Then, to examine whether this proliferative effect by IL-13 is mediated by IL-13Rα2, we applied a knockdown system using commercially available siRNA to MO-LAS-B cells. Efficient knockdown of *IL13RA2* was obtained at the mRNA level by specific siRNA (Fig. S1). *IL13RA2* knockdown decreases cell proliferation under the IL-13 supplementation (Fig. 2G). Furthermore, the addition of anti-IL-13 neutralizing antibodies made the difference less noticeable between control siRNA-transfected cells and *IL13RA2* siRNA-transfected cells in the presence of IL-13, implicating the involvement of IL-13 in cell proliferation (Fig. 2G). These findings highlight the critical role of the IL-13-IL-13Rα2 axis in angiosarcoma cell proliferation.

**Fig 2.**
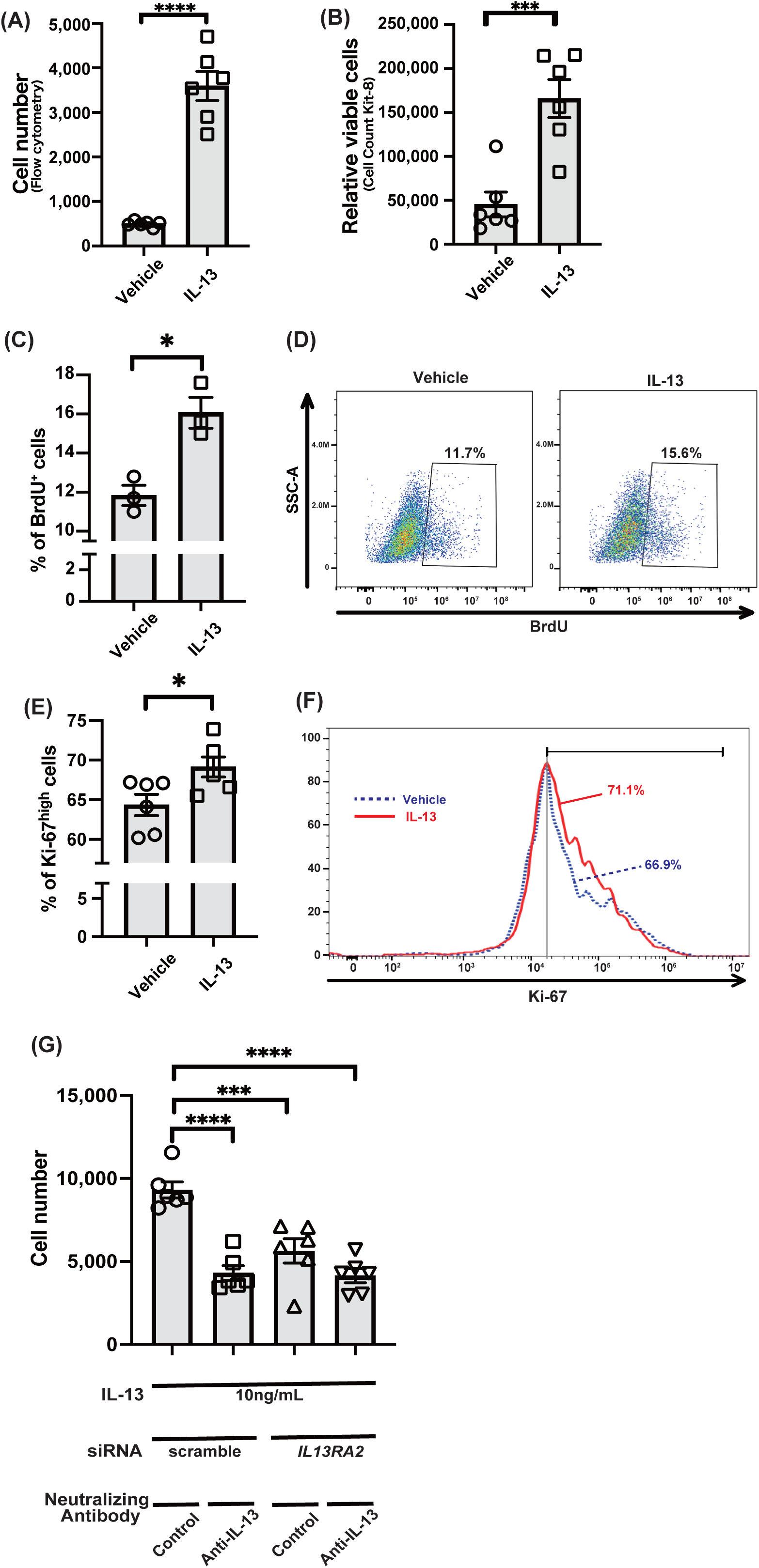
Accelerated proliferation in angiosarcoma cells mediated by IL-13/IL-13Rα2 axis. (A, B) The effect of IL-13 (10 ng/mL) for 24 hours on MO-LAS-B cell numbers was evaluated by flow cytometry (A) and Cell Counting Kit-8 cell proliferation assay (B) (*n* = 6 for each group). (C) Percent of bromodeoxyuridine (BrdU)-positive cells after vehicle or IL-13 (10 ng/mL) treatment for 48 hours (*n* = 3 for each group). (D) Representative flow cytometric analysis of BrdU-positive cells from the vehicle or IL-13 treated cells. Numbers show percentages of cells in the indicated gate. (E) Percent of Ki-67^hi^ cells after vehicle or IL-13 (10 ng/mL) treatment for 48 hours (*n* = 6 for each group). (F) Representative histograms from Ki-67 staining of angiosarcoma cells, MO-LAS-B. Percent of Ki-67high cells in each sample was indicated. Two histograms in this figure were representative of 6 samples in each group. (G) Cell proliferation assay demonstrating the effects of IL-13-IL13Rα2 axis on the MO-LAS-B cell numbers. All populations were treated with IL-13 (10 ng/mL) for 72 hours. Some samples were transfected with scrambled siRNAs or *IL13RA2* siRNAs 48 hours before IL-13 stimulation. Some samples were cultured with anti-IL-13 neutralizing antibodies 1 hour before IL-13 stimulation (*n* = 6 for each group). (A, B, C, E, G) Representative data from one of at least two independent experiments are shown. \**p*<0.05, \*\*\**p*<0.001, \*\*\*\**p*<0.0001.

### Potential positive feedback loop mechanism in IL-13-stimulated angiosarcoma cells

We additionally investigated the effects of IL-13 on the expression of IL-13Rα2. IL-13 stimulation increased IL-13Rα2 expression (Fig. 3A), suggesting a positive feedback loop between IL-13 and IL-13Rα2. Given the reported involvement of STAT6 in the downstream pathway of IL-13Rα2^25^, we further treated cells with a STAT6 inhibitor, AS1517499, which abolished the positive feedback loop (Fig. 3A). These findings implicate that IL-13 signaling through IL-13Rα2 in angiosarcoma cells is mediated by STAT6 activation. Considering the reported critical role of VEGF-A in tumor angiogenesis and its correlation with angiosarcoma cell proliferation^8^, we confirmed that VEGF-A supplementation promotes MO-LAS-B cell proliferation (Fig. 3B). Then, we examined the effect of IL-13 on *VEGFA* expression and found that IL-13 stimulation significantly upregulated *VEGFA* mRNA expression, which was similarly suppressed by AS1517499 (Fig. 3C). These results raise the possibility that increased production of VEGF-A is involved in cell proliferation induced by IL-13.

**Fig 3.**
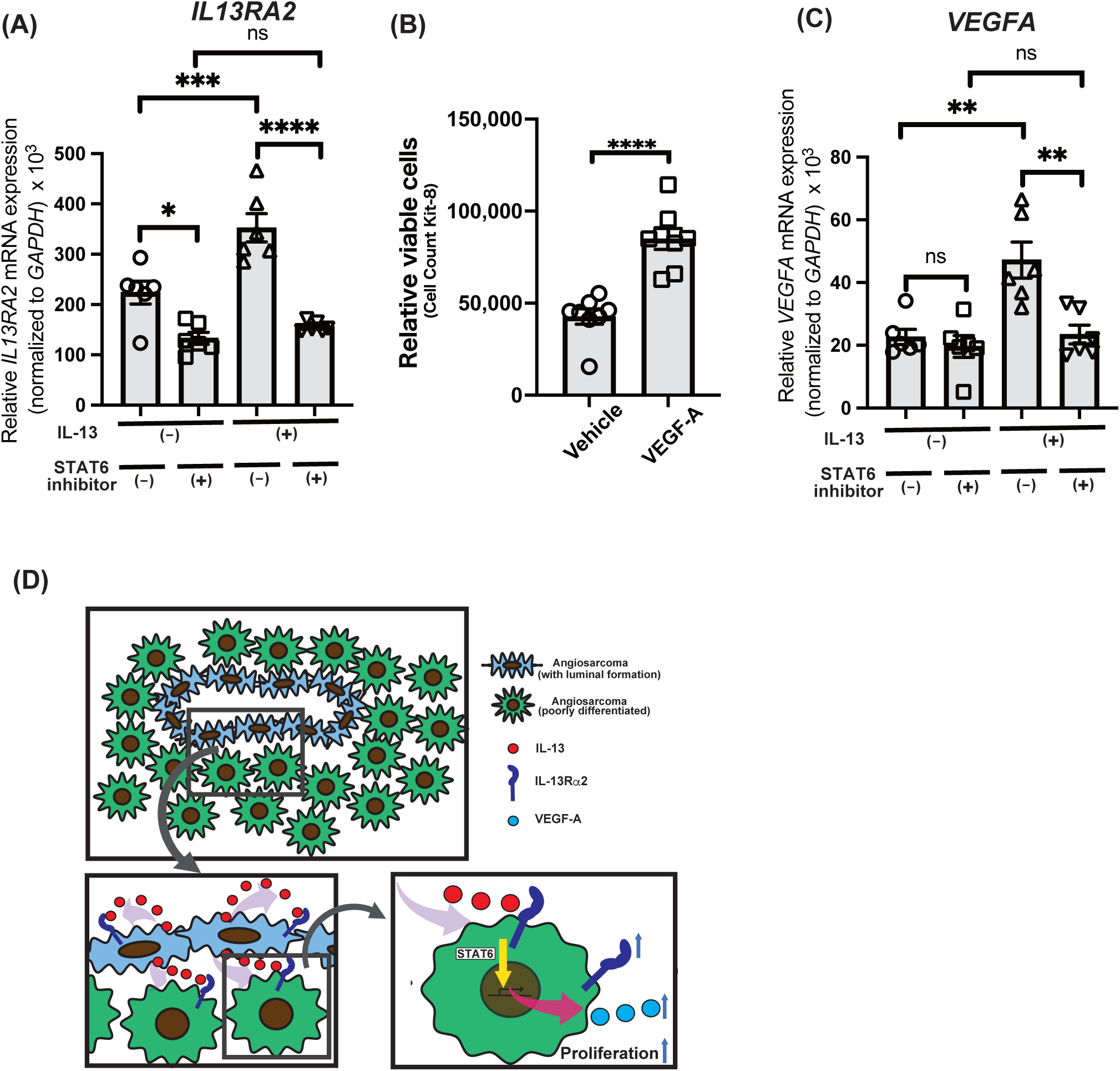
IL-13-induced upregulated expression of *IL13RA2* and *VEGFA* via STAT6 activation. (A, C) Relative mRNA expression of *IL13RA2* (A) and *VEGFA* (C) under vehicle or IL-13 treatment (10 ng/mL). Some samples were treated with STAT6 inhibitor, AS1517499, 1 hour before the vehicle or IL-13 stimulation, (*n* = 6 for each group). (B) CCK-8 cell proliferation assay evaluating the effect of VEGF-A on MO-LAS-B cell proliferation, MO-LAS-B cells were stimulated with vehicle or VEGF-A (50ng/ml) for 48 hours (*n* = 8 for each group). (D) Conceptual model for the role of IL-13 in the proliferation of angiosarcoma cells. IL-13Rα2 is highly expressed in angiosarcoma, and IL-13 promotes proliferation through IL-13Rα2. Moreover, IL-13Rα2 and VEGF expression are upregulated by IL-13 stimulation in a STAT6-dependent manner. (A-C) Representative data from one of at least two independent experiments are shown. \*\**p*<0.01, \*\*\**p*<0.001, \*\*\*\**p*<0.0001.

## Discussion

This study investigated the role of IL-13 and its receptors in angiosarcoma pathogenesis. Our findings revealed that IL-13 is present in the angiosarcoma microenvironment, with strong expression observed in atypical tumor cells. Notably, angiosarcoma cell lines exhibited higher expression of IL-13Rα2 compared to other cell types, including immune cells and skin fibroblasts. This observation was further confirmed in patient-derived angiosarcoma samples through immunohistochemistry. Functional studies showed that IL-13 promotes angiosarcoma cell proliferation, an effect mediated primarily through IL-13Rα2. Knockdown experiments and neutralizing antibody studies corroborated the critical role of the IL-13-IL-13Rα2 axis in driving angiosarcoma cell proliferation. Furthermore, IL-13 stimulation increased IL-13Rα2 and VEGF-A expression, suggesting a positive feedback mechanism. This upregulation was attenuated by a STAT6 inhibitor, suggesting a potential role for STAT6 in this signaling pathway. These findings underscore the importance of the IL-13-IL-13Rα2 pathway in angiosarcoma progression and highlight its potential as a therapeutic target (Fig. 3D).

In the immunohistochemical staining for IL-13, atypical cells forming luminal structures showed strong positivity. Of note, not all cells were uniformly stained, potentially reflecting varying degrees of cellular differentiation. Previous studies have reported that IL-13 can exert effects even at low concentrations and demonstrate concentration-dependent actions^28^. Therefore, even in the absence of robust staining signals, IL-13 may still be functionally active if its receptors are expressed. Conversely, IL13Rα2 exhibited diffuse staining patterns across tumor cells, suggesting widespread expression in the neoplasm. These findings indicate that even within the same angiosarcoma, the roles and mechanisms of IL-13 and its receptor may differ based on factors such as differentiation status.

This study showed remarkably high expression of IL-13Rα2 in the angiosarcoma cell line MO-LAS-B. Immunohistochemical analysis corroborated the overexpression of IL-13Rα2 in angiosarcoma specimens compared to hemangiomas. In contrast, IL-13Rα1 expression was limited in the angiosarcoma cell line, indicating a distinct pattern of high IL-13Rα2 and low IL-13Rα1 expression in angiosarcoma. Our investigations further demonstrated that IL-13 stimulation induces IL-13Rα2 expression. These observed IL-13 receptor expression patterns suggest that IL-13 signaling is mediated predominantly through IL-13Rα2, potentially indicating a positive feedback mechanism in which the IL-13-IL-13Rα2 axis augments IL-13Rα2 expression. Although the mechanisms of IL-13Rα2 expression in other malignancies remain to be fully understood, our findings offer insights into one aspect of IL-13Rα2 expression regulation. A key limitation in validating this hypothesis is the lack of suitable *in vitro* models due to the undeveloped and difficult-to-obtain IL-13Rα2 specific inhibitors.

While our findings demonstrate that IL-13 promotes angiosarcoma proliferation, the precise underlying mechanisms remain elusive. We have shown that IL-13 upregulates VEGF-A expression via STAT6 activation, possibly partially explaining the proliferative effect. However, questions persist about whether IL-13 acts directly or induces other growth factors through IL-13Rα2 signaling, potentially involving a combination of these processes. Given the multitude of potential pathways involved, further investigation is necessary to identify additional mechanisms in IL-13-mediated angiosarcoma progression.

The present study integrated expression analysis of clinical specimens with *in vitro* investigations using cell lines. These approaches provided valuable insights but have inherent limitations. The lack of a well-established animal model for angiosarcoma necessitated our experimental design. Notably, our *in vitro* results may not fully reflect *in vivo* conditions. Furthermore, our *in vitro* experiments exclusively utilized MO-LAS-B cells, potentially limiting the representation of complex cellular interactions within the skin microenvironment. Future development of advanced animal models or co-culture systems would facilitate more robust validation of our findings. Moreover, our study focused on cutaneous angiosarcoma. Thus, angiosarcomas arising in other anatomical locations may display distinct biological characteristics. Further investigations examining angiosarcomas from various anatomical sites are essential to determine the broader applicability of our results.

Chimeric antigen receptor (CAR)-T cell therapy targeting IL13Rα2 has demonstrated potential in treating glioblastoma and other IL13Rα2-positive cancers^29^. Clinical studies have shown tumor regression in glioblastoma patients treated with IL13Rα2-targeted CAR-T cells^30^. However, challenges remain, such as potential cross-reactivity with IL13Rα1^31^ and the need for improved persistence and efficacy. Recent advancements include combination therapies with checkpoint inhibitors^32^, which have shown enhanced anti-tumor activity in preclinical models. These approaches may lead to improved outcomes for patients with IL13Rα2-positive tumors. Our findings implicate that CAR-T cell therapy may be a viable treatment option for angiosarcoma.

Current therapeutic targets for angiosarcoma primarily focus on pathways such as VEGFR1-3, mTOR, MEK, c-Myc, PD-1/PD-L1, and β-adrenergic receptors^33^. Cytokine-targeted therapies, however, are limited, with interleukin-2 (IL-2) being the main immunotherapeutic option explored. IL-2 therapy shows limited efficacy and high toxicity when administered systemically, and while local administration methods are being researched, their effectiveness in angiosarcoma remains unclear^34^. Our findings suggest a promising cytokine-targeted approach for angiosarcoma. This novel mechanism could potentially be integrated with current treatments, possibly leading to improved therapeutic outcomes and patient quality of life.

## Materials and methods

### Patients and clinical samples

The study protocol conforms to the ethical guidelines of the 1975 Declaration of Helsinki. It was approved by the Institutional Research Ethics Committee of the Faculty of Medicine of The University of Tokyo. Informed consent was obtained from the patients for the use of all samples. Skin samples were obtained from patients with angiosarcoma, hemangioma, and atopic dermatitis. Control skin samples were obtained from healthy donors.

### Cell cultures

Two types of human angiosarcoma cell lines, MO-LAS-B and ISO-HAS-B cells, were obtained from the Cell Resource Center for Biomedical Research, Institute of Development, Aging and Cancer (Tohoku University, Sendai, Japan). MO-LAS-B cells was originally derived from a patient with metastatic scalp angiosarcoma to the pleura, while ISO-HAS-B cells was established from a patient with primary scalp angiosarcoma^35,36^. These cell lines were cultured in Dulbecco’s Modified Eagle’s Medium (DMEM; Fujifilm Wako Pure Chemical Corporation, Osaka, Japan) supplemented with 10% fetal bovine serum (FBS) and 1% penicillin-streptomycin (100 U/mL penicillin and 100 mg/mL streptomycin). Given the lost cell viability under the complete serum starvation in MO-LAS-B cells, we cultured the cells in a serum-reduced (5% FBS) medium for the indicated time before stimulation. Human skin squamous cell carcinoma line HSC5 and epidermoid carcinoma line A431 were obtained from the Japanese Collection of Research Bioresources Cell Bank (Osaka, Japan)^37^. HSC5 and A431 cells were maintained in DMEM supplemented with 10% FBS^38^. Regarding human dermal skin fibroblasts, clinically obtained skin samples were cut into small pieces and placed on Corning 24-well plates. Then, the dermis pieces were removed a few days after the fibroblast growth on the dish was checked. Dermal fibroblasts were cultured in DMEM supplemented with 10% FBS and antibiotics. Cells were incubated at 37°C in a humidified 5% CO_2_ atmosphere.

### Cell sorting

Peripheral blood mononuclear cells (PBMCs) were isolated from EDTA-anticoagulated whole blood using 1X RBC Lysis Buffer (Thermo Fisher Scientific, Waltham, MA, USA) according to the manufacturer’s instructions. PBMCs were stained with fluorochrome-conjugated monoclonal antibodies against human CD45 (PE/Cy7), CD19 (PerCP/Cy5.5), CD3 (FITC), CD14 (PE), CD16 (APC) (all from BioLegend, San Diego, CA, USA), and Fixable Viability Dye eFluor 780 (Thermo Fisher Scientific) for 30 min at 4°C. Stained cells were sorted into neutrophils (CD45^+^CD16^+^), T cells (CD45^+^CD3^+^), and B cells (CD45^+^CD19^+^) (gated on the basis of forward and side scatter profiles) using a Sony SH800S Cell Sorter (Sony Biotechnology Inc., San Jose, CA, USA).

### Gene expression analysis

Gene expression in sorted cells and cell lines was analyzed using the QPCR method. RNA isolation and QPCR analysis were performed as described previously^39^. Briefly, the RNeasy Mini Kit (Qiagen, Crawley, UK) extracted total RNA from cells according to the manufacturer’s instructions. The extracted RNA was then reverse-transcribed to cDNA using the ReverTraAce qPCR RT Master Mix (Toyobo, Osaka, Japan). Quantitative analyses of mRNA expression levels were conducted using specific primers designed by Primer 3 online software. mRNA expression levels were evaluated using SYBR Green qPCR Master Mix (TOYOBO, Osaka, Japan). The reactions were performed on a StepOne Real-Time PCR System (Applied Biosystems, Foster City, CA, USA). *GAPDH* was used as an internal control to normalize the amounts of loaded cDNA. Relative gene expression was calculated using the comparative ΔΔCT method. The sequences of the primers used in this study are provided in Table S1.

### Histological and immunohistochemical analyses

Deparaffinized sections (3-μm thick) were stained for IL-13RA2, as described previously^40,41^. Frozen sections (6-μm thick) were prepared for IL-13 by embedding samples in OCT compound (Sakura Finetek, Japan). The following antibodies were used: IL-13RA2/CD213a2 (E7U7B) Rabbit mAb (#85677, Cell Signaling Technology, Danvers, MA, USA) and purified anti-human IL-13 antibody (#501902, BioLegend, San Diego, CA, USA). Tissues were subsequently stained with an avidin-biotin-peroxidase complex using R.T.U. VECTASTAIN Universal Elite ABC Kit (PK-7200, Vector Laboratories, Burlingame, CA, USA). All analyses were performed using a Keyence BZ-X800 fluorescence microscope (Keyence Corporation, Osaka, Japan).

### Flow cytometry analysis

Flow cytometry was performed to analyze cell surface expression and evaluate cell proliferation. Cells were harvested and washed twice with FACS buffer (PBS containing 2% FBS and 2 mM EDTA). Dead cells were excluded by staining with Fixable Viability Dye eFluor 780 (Thermo Fisher Scientific). For IL-13Rα2 analysis, cells were incubated with human IL-13 R alpha 2 antibody (AF146, R&D Systems, Minneapolis, MN, USA) followed by donkey anti-goat IgG (H+L) cross-adsorbed secondary antibody, Alexa Fluor 488 (A-11055, Invitrogen, Carlsbad, CA, USA) for 30 minutes at 4°C. For cell proliferation analysis, cells were fixed and permeabilized using BD Cytofix/Cytoperm Fixation/Permeabilization Solution Kit (554714, BD Biosciences, San Jose, CA, USA) according to the manufacturer’s instructions, then stained with monoclonal mouse anti-human Ki-67 antigen, clone MIB-1 (M724029, DAKO, Glostrup, Denmark) followed by PE goat anti-mouse IgG (minimal x-reactivity) (405307, BioLegend). Samples were analyzed using a CytoFLEX S flow cytometer (Beckman Coulter, Brea, CA, USA) and data were processed using FlowJo software (v10.6.1, TreeStar, Ashland, OR, USA). A minimum of 10,000 events were acquired for each sample.

### Bromodeoxyuridine (BrdU) labeling and cell stimulation

Following serum-reduced starvation (5% FBS) for 12 hours in DMEM, MO-LAS-B cells were stimulated with recombinant human IL-13 (200-13, PeproTech, Rocky Hill, NJ, USA) at 10 ng/mL or left unstimulated (control). After 24 hours of stimulation, BrdU (B5002, Sigma-Aldrich, St. Louis, MO, USA) was added to the culture medium at a final concentration of 10 μM. Cells were then incubated with BrdU for an additional 24 hours. Staining for flow cytometry analyses was performed according to the BD Pharmingen BrdU Flow Kit protocol (557891, BD Biosciences, San Jose, CA, USA) as previously used^42^ Briefly, cells were fixed and permeabilized using BD Cytofix/Cytoperm Buffer, treated with DNase I (D4513-1VL, Sigma-Aldrich, St. Louis, MO, USA) to expose incorporated BrdU, and stained with APC-conjugated anti-BrdU antibody (included in the kit).

### Small interfering RNA (siRNA) and transfection

MO-LAS-B cells were grown to 40% confluence in 10cm-dishes before transfection with Silencer Select Human *IL13RA2* siRNA (s7375, Thermo Fisher Scientific, Waltham, MA, USA) or Silencer Select Negative Control No.1 siRNA (4390843, Thermo Fisher Scientific) using Lipofectamine 3000 Transfection Reagent (L3000001, Thermo Fisher Scientific). The final siRNA concentration was 40 nM. Transfection efficiency was assessed by QPCR and flow cytometry analysis 48 hours post-transfection.

### Cell counting

Cell number was assessed using two methods: flow cytometry and CCK-8 assay. Flow cytometry: Cells were seeded in 12-well plates. After indicated treatments, cells were harvested and analyzed by flow cytometry as described in the Flow cytometry analysis section. The total number of viable cells was determined based on forward and side scatter properties and viability dye exclusion. CCK-8 assay: Cells were seeded in 12-well plates. After indicated treatments, cell proliferation was measured using the Cell Counting Kit-8 (CCK-8, Dojindo Molecular Technologies, Kumamoto, Japan) according to the manufacturer’s instructions. Absorbance was measured at 450 nm using a microplate reader (SpectraMax i3x, Molecular Devices, San Jose, CA, USA).

### IL-13 treatment and neutralizing antibody treatment

Before IL-13 stimulation, cells were subjected to serum starvation for 12 hours in 5% FBS to synchronize cell cycles and minimize the effects of serum-derived growth factors. Following starvation, cells were treated with recombinant human IL-13 (200-13, PeproTech, Rocky Hill, NJ, USA) at 10 ng/mL for indicated times. Vehicle-treated cells served as controls. For neutralization experiments, a mixture containing 0.8 mg/mL of anti-IL-13 neutralizing antibody (clone JES10-5A2, BioLegend) and recombinant IL-13 at a final concentration of 10 ng/mL was prepared in 0.1% BSA DMEM. This mixture was pre-incubated for 30 minutes at room temperature before adding to the cells in 12-well plates. Cells were then cultured in the presence of this neutralization mixture for the indicated amount of time.

### STAT6 inhibition

Cells were pre-treated with the STAT6 inhibitor AS1517499 (500 nM, Axon Medchem, catalog Axon 1992) or vehicle control (0.1% DMSO) for 1 hour before stimulation with IL-13 (10 ng/mL). The inhibitor was maintained in the culture medium throughout the experiment to ensure continuous STAT6 inhibition. Cells were harvested 24 hours after IL-13 stimulation for subsequent analysis.

### Statistical analysis

Prism (GraphPad, ver. 10) software was used for all statistical analyses. Statistical analysis was performed by one-way ANOVA with Dunnett’s post hoc tests for multiple group comparisons and the two-tailed statistical analysis. Two-tailed, unpaired *t*-tests were performed when comparing only two groups. A value of P < 0.05 was considered statistically significant. In graphs, horizontal lines indicate means and error bars indicate standard error of the mean (SEM).

## Data availability

The original data analyzed in this study are available from the corresponding author upon reasonable request.

## Acknowledgments

We would like to thank the medical staff and patients in the Department of Dermatology at The University of Tokyo Hospital for contributing to this study.

## Author contributions

H.S. [Hayakazu Sumida] designed the study, analyzed the results, wrote the manuscript, and discussed the results. H.S. [Hinako Saito] conducted most of the experiments, analyzed the results, and wrote the manuscript. S.S. discussed the results and supervised experiments. I.O., O.I., A.S., H.N. and R.I. were involved in data curation and revising the manuscript. All authors reviewed and approved the final manuscript.

## Funding

JSPS KAKENHI Grant Number 24K02469 and 23K18286 [to HS(Hayakazu Sumida)] from the Japan Society for the Promotion of Science (JSPS).

## Competing interests

The authors declare no competing interests.

## Notes

### Competing Interest Statement

The authors have declared no competing interest.

