## Supplemental Figure 1 and Table 1 for "IL-13/IL-13Rα2 axis promotes proliferation of angiosarcoma cells"

#### Supplementary information

### Supplementary Figure 1

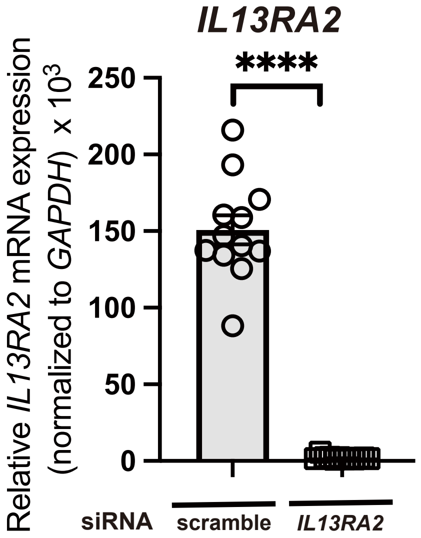


**Supplementary Figure 1.**

**Knockdown efficiency of IL13RA2 siRNA in MO-LAS-B cells.**

Relative mRNA expression of IL13RA2 in MO-LAS-B cells transfected with scramble or IL13RA2-specific siRNAs, showing more than 95% knockdown efficiency in IL13RA2 mRNA expression with specific siRNA ($n = 12$ for each group).

The analysis confirmed >95% knockdown efficiency. ****$p<0.0001$.

**Supplementary information**

**Supplementary Table 1.**

**List of primers used for real-time PCR.**

**Human**

| Gene | Forward Primer (5'-3') | Reverse Primer (5'-3') |
|---|---|---|
| *IL13RA2* | GGAGCATACCTTTGGGACCTATTC | ATTGTCGGGTTTCATTTGTTGTTT |
| *IL13RA1* | GGGAAAAGGGAGGGAAAAGG | TGACACTGGGTTTGCTTAGGTATG |
| *VEGFA* | AAGGAGGAGGGCAGAATCAT | TGGTGATGTTGGACTCCTCA |
| *GAPDH* | CACCCACTCCTCCACCTTTG | CTCTCTTCCTCTTGTGCTCTTGCT |
